# Genome-wide association meta-analysis of PR interval identifies 47 novel loci associated with atrial and atrioventricular electrical activity

**DOI:** 10.1101/241489

**Authors:** Jessica van Setten, Jennifer A. Brody, Yalda Jamshidi, Brenton R. Swenson, Anne M. Butler, Harry Campbell, M. Fabiola Del Greco, Daniel S. Evans, Quince Gibson, Daniel F. Gudbjartsson, Kathleen F. Kerr, Bouwe P. Krijthe, Leo-Pekka Lyytikäinen, Christian Müller, Martina Müller-Nurasyid, Ilja M. Nolte, Sandosh Padmanabhan, Marylyn D. Ritchie, Antonietta Robino, Albert V. Smith, Maristella Steri, Toshiko Tanaka, Alexander Teumer, Stella Trompet, Sheila Ulivi, Niek Verweij, Xiaoyan Yin, David O. Arnar, Folkert W. Asselbergs, Joel S. Bader, John Barnard, Josh Bis, Stefan Blankenberg, Eric Boerwinkle, Yuki Bradford, Brendan M. Buckley, Mina K. Chung, Dana Crawford, Marcel den Hoed, Josh Denny, Anna F. Dominiczak, Georg B. Ehret, Mark Eijgelsheim, Patrick T. Ellinor, Stephan B. Felix, Oscar H. Franco, Lude Franke, Tamara B. Harris, Hilma Holm, Gandin Ilaria, Annamaria Iorio, Mika Kähönen, Ivana Kolcic, Jan A. Kors, Edward G. Lakatta, Lenore J. Launer, Honghuang Lin, Henry J. Lin, Ruth J.F. Loos, Steven A. Lubitz, Peter W. Macfarlane, Jared W. Magnani, Irene Mateo Leach, Thomas Meitinger, Braxton D. Mitchell, Thomas Munzel, George J. Papanicolaou, Annette Peters, Arne Pfeufer, Peter P. Pramstaller, Olli T. Raitakari, Jerome I. Rotter, Igor Rudan, Nilesh J. Samani, David Schlessinger, Claudia T. Silva Aldana, Moritz F. Sinner, Jonathan D. Smith, Harold Snieder, Elsayed Z. Soliman, Timothy D. Spector, David J. Stott, Konstantin Strauch, Kirill V. Tarasov, Andre G. Uitterlinden, David R. van Wagoner, Uwe Völker, Henry Völzke, Melanie Waldenberger, Harm Jan Westra, Philipp S. Wild, Tanja Zeller, Alvaro Alonso, Christy L. Avery, Stefania Bandinelli, Emelia J. Benjamin, Francesco Cucca, Marcus Dörr, Luigi Ferrucci, Paolo Gasparini, Vilmundur Gudnason, Caroline Hayward, Susan R. Heckbert, Andrew A. Hicks, J. Wouter Jukema, Stefan Kääb, Terho Lehtimäki, Yongmei Liu, Patricia B. Munroe, Afshin Parsa, Ozren Polasek, Bruce M. Psaty, Dan M. Roden, Renate B. Schnabel, Gianfranco Sinagra, Kari Stefansson, Bruno H. Stricker, Pim van der Harst, Cornelia M. van Duijn, James F. Wilson, Sina Gharib, Paul I.W. de Bakker, Aaron Isaacs, Dan E. Arking, Nona Sotoodehnia

## Abstract

Electrocardiographic PR interval measures atrial and atrioventricular depolarization and conduction, and abnormal PR interval is a risk factor for atrial fibrillation and heart block. We performed a genome-wide association study in over 92,000 individuals of European descent and identified 44 loci associated with PR interval (34 novel). Examination of the 44 loci revealed known and novel biological processes involved in cardiac atrial electrical activity, and genes in these loci were highly over-represented in several cardiac disease processes. Nearly half of the 61 independent index variants in the 44 loci were associated with atrial or blood transcript expression levels, or were in high linkage disequilibrium with one or more missense variants. Cardiac regulatory regions of the genome as measured by cardiac DNA hypersensitivity sites were enriched for variants associated with PR interval, compared to non-cardiac regulatory regions. Joint analyses combining PR interval with heart rate, QRS interval, and atrial fibrillation identified additional new pleiotropic loci. The majority of associations discovered in European-descent populations were also present in African-American populations. Meta-analysis examining over 105,000 individuals of African and European descent identified additional novel PR loci. These additional analyses identified another 13 novel loci. Together, these findings underscore the power of GWAS to extend knowledge of the molecular underpinnings of clinical processes.

## Introduction

The PR interval on the surface electrocardiogram reflects atrial and atrioventricular node myocyte depolarization and conduction. Abnormalities in PR interval duration are associated with increased risk of atrial fibrillation, which carries substantial risk of morbidity and mortality, and with cardiac conduction defects and heart block, conditions that often necessitate pacemaker implantation.^1^ Understanding the molecular mechanisms underlying PR interval may provide insights into cardiac electrical disease processes, and identify potential drug targets for prevention and treatment of atrial fibrillation and conduction disease.

Twin and family studies suggest that the heritability of PR interval is between 40% and 60%.^2^ Prior genome-wide association studies (GWAS) in up to 30,000 individuals have identified ten loci associated with PR interval among European-descent individuals.^3,4^ The key motivation for the present study was to extend the biological and clinical insights derived from GWAS data by utilizing a >3 fold increased sample size to detect novel PR loci. We further increased power by performing trans-ethnic meta-analyses. To gain additional biological and clinical insights, we tested for pleiotropy with other clinically relevant phenotypes. We examined the biological and functional relevance of identified associations to gain insights into molecular processes underlying clinically important outcomes.

### Meta-analysis of Genome-Wide Association Studies for PR interval among European Ancestry Individuals

We meta-analyzed ~2.7 million single nucleotide polymorphisms (SNPs) from GWAS data on approximately 92,340 individuals of European ancestry from 31 studies (**Supplementary Table 1a and 1b)** for association with PR interval using an additive genetic model. A total of 1,601 SNPs mapping to 44 loci (of which 34 novel in Europeans) reached genome-wide significance (*P* ≤ 5 × 10^−8^) (Figure 1, Table 1, **Supplementary Figures 1 and 2**). While genomic inflation factor lambda was modest (1.11), linkage disequilibrium (LD) score regression^5^ showed that deviation from the expected P-value distribution was mainly caused by true polygenicity (**Supplementary Figure 3**). Using a Bayesian gene-based test of association (GWiS),^6^ we identified 61 independent signals in the 44 loci. For example, the top locus on chromosome 3, mapping to the two cardiac sodium channel genes *SCN5A* and *SCN10A,* had seven independent signals associated with PR interval (Figure 2a).

**Figure 1:**
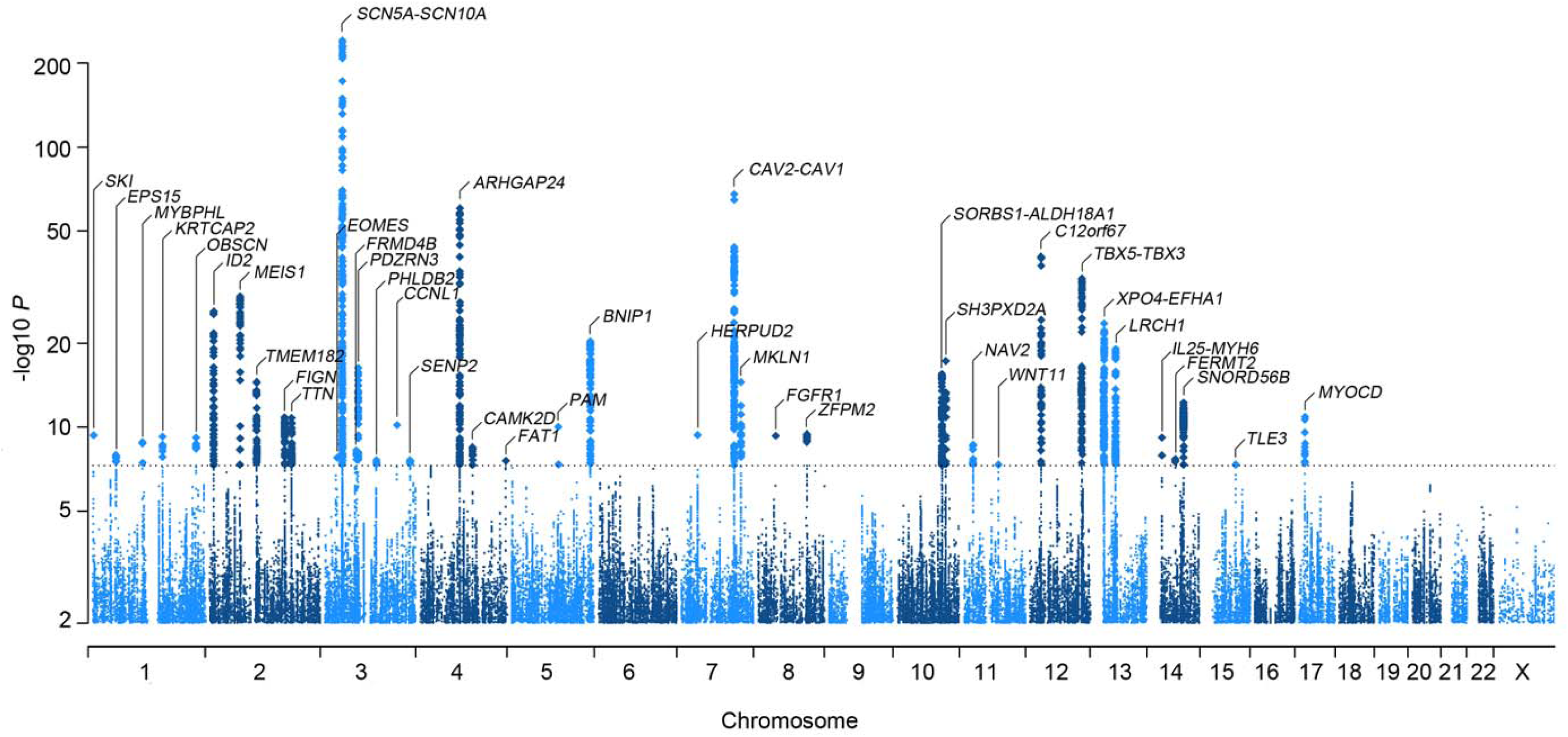
Genome-wide results of PR interval in 92,000 individuals of European descent. 2.8 million SNPs were tested association with PR interval in 31 cohorts. The Manhattan plot shows the meta-analysis association results: 44 independent loci (labeled) are associated at the genome-wide significance level of *P* ≤ 5 × 10^−8^, as marked by the dashed line.

**Table 1:**
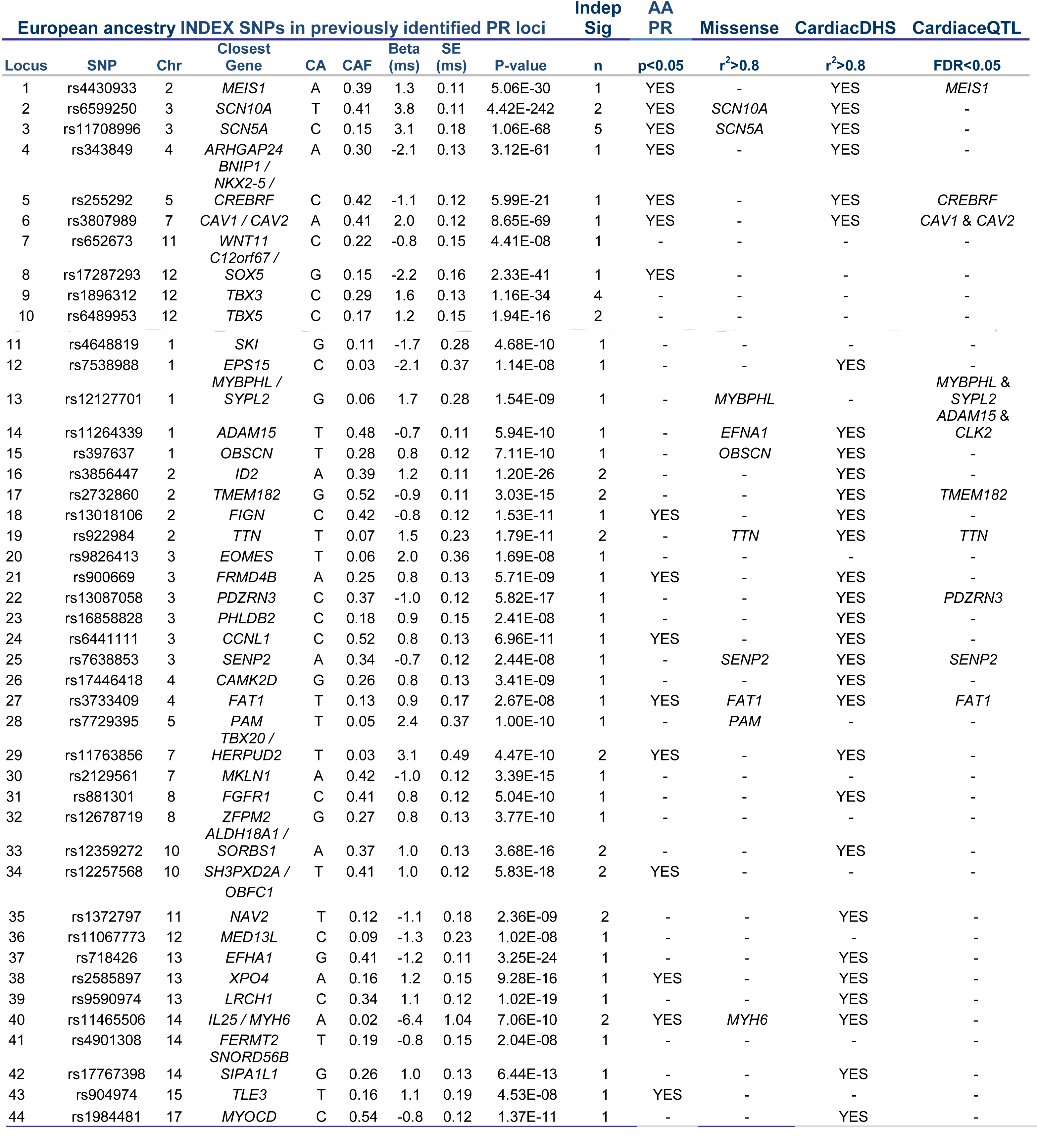
Description of novel and previously identified loci. For each locus we list the number of independent signals, whether this locus is nominal significant in African Americans, if missense SNPs are in LD with the index SNP or SNPs, if the index SNP is in LD with or located in a cardiac DHS, and if the locus contains cardiac or blood eQTLs. Abbreviations: Chr - chromosome, CA - coded allele, CAF - coded allele frequency, SE - standard error.

**Figures 2a-c:**
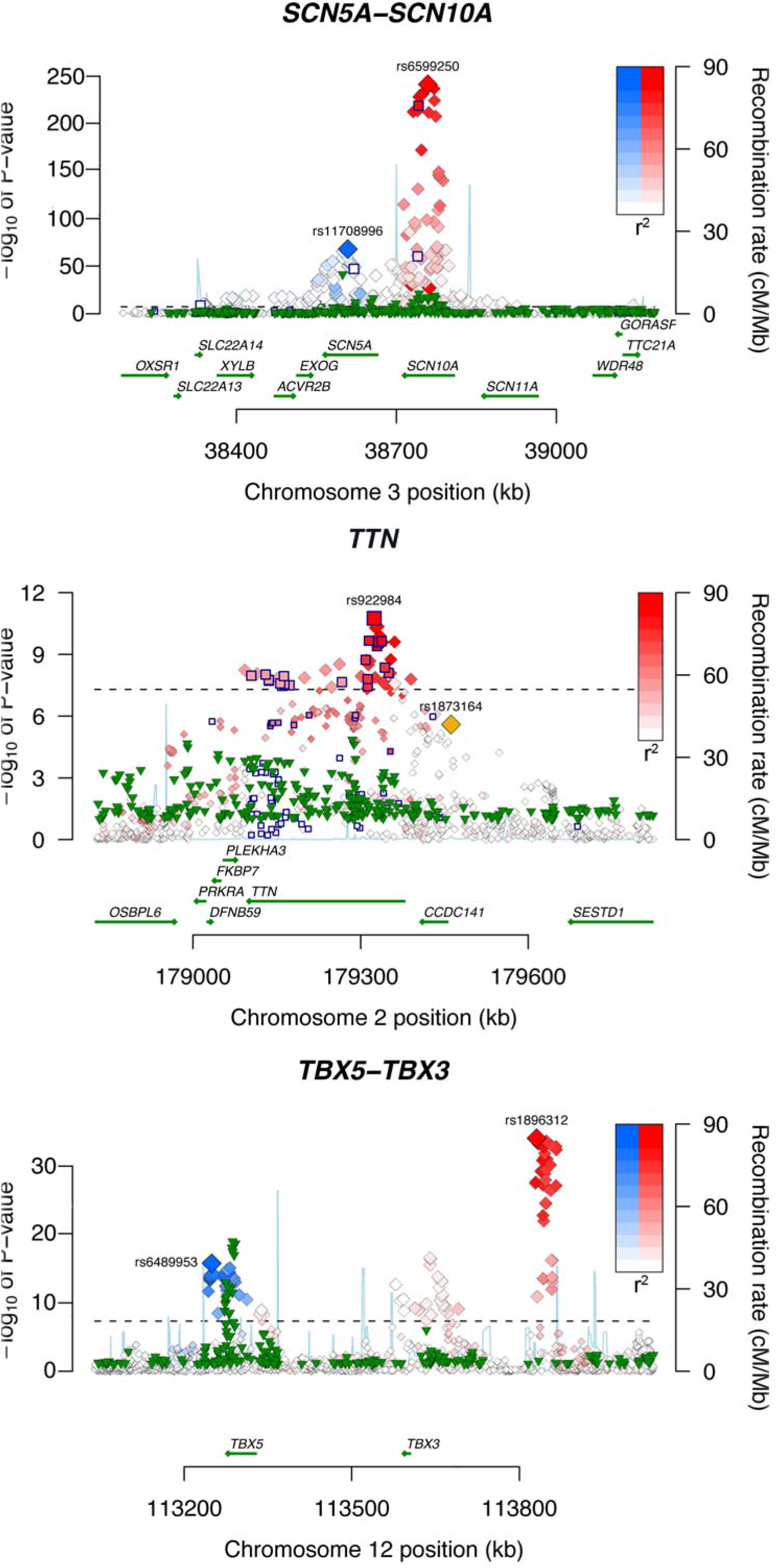
Regional association plots of specific loci associated with PR interval. Each SNP is plotted with respect to its chromosomal location (x axis) and its P value (y axis on the left). The blue line indicates the recombination rate (y axis on the right) at that region of the chromosome. Blue outlined squares mark non-synonymous SNPs. Green triangles depict association results of the African Americans meta-analysis, only SNPs with *P* < 0.1 are shown. (a) Locus 2 and 3 (*SCN10A*-*SCN5A*) on chromosome 3. The index SNPs for the two genes are named with their rs-numbers and highlighted with two different colors (blue and red). Other SNPs in linkage disequilibrium with the index SNP are denoted in the same color; color saturation indicates the degree of correlation with the index SNP. (b) Locus 19 (*TTN*) on chromosome 2; and (c) Locus 9 and 10 (*TBX5*-*TBX3*) on chromosome 12.

### Putative Functional Variants

To assess the functional relevance of the identified SNPs, we examined whether the index variants were in high LD with either nonsynonymous variants or with putative regulatory SNPs. Ten of the 44 loci had missense variants in high LD (r^2^ > 0.8) with the index SNP (Table 1, **Supplementary Table 3)**. *TTN,* in particular, was enriched for missense SNPs, with the top signal and approximately one-third of the 47 genome-wide significant TTN SNPs being missense (Figure 2b). To examine the possible impact of these amino acid substitutions on protein structure or function, we used two prediction algorithms, Sift^7^ and PolyPhen-2^8^. The vast majority of the genome-wide significant missense variants at the 44 loci were categorized as tolerated by Sift and benign by PolyPhen-2, consistent with modest effects on PR interval not subjected to purifying selection (**Supplementary Table 3)**.

Expression quantitative trait locus (eQTL) analysis suggests that index SNPs in half of the identified loci (22/44) are involved in *cis* gene regulation in at least one of the two tissue types examined at a false discovery rate (FDR) of <0.05 (**Supplementary Table 4**): left atrial appendage (n = 230 samples, 10 eQTL SNPs) and whole blood (n = 5311 samples, 16 eQTL SNPs). Several points are worth highlighting. First, for most of the 22 loci, the eQTL associations are for the gene nearest the index SNP, but for nearly one-third, they are not. Second, certain SNPs can be promiscuous in that they are associated with the transcript expression of multiple different genes. Third, despite substantially greater power to detect associations in whole blood compared to cardiac tissue due to markedly larger sample size, most of the eQTL associations found in cardiac atrial tissue – e.g. associations with *MEIS1* (**Supplementary Figure 4a**), *CAV1*, *FAT1*, and *TTN* transcripts – were not found in whole blood samples and appear to have some tissue-specificity. Two eQTL associations were found in both blood and cardiac tissue (*ADAM15* and *SENP2* (**Supplementary Figure 4b**). While several other index SNPs were also associated with eQTLs in both tissue types, they were associated with transcript expression levels of different genes. For instance, locus 17 SNP rs2732860 was associated with *TMEM182* expression in atrial tissue but with *MFSD9* expression in blood, again suggesting tissue-specificity for SNP–eQTL associations. Taken together, these data underscore the importance of examining eQTL data in tissue types relevant to the trait of interest: even with a modest study size of 230 cardiac atrial samples, a notable number of eQTL associations were uncovered.

The majority of loci (30/44) contain index SNPs that lie in, or are in high LD with, regulatory regions of the genome that are marked by deoxyribonuclease I (DNAse I) hypersensitivity sites (DHSs), lending further support to the hypothesis that regulation of gene-expression plays an important role in determining PR interval (Table 1). To provide insight into the cellular and tissue structure of the phenotype, we examined *P*-values of all SNPs in the PR meta-analysis and assessed cell and tissue selective enrichment patterns of progressively more strongly associated variants to explore localization of signal within specific lineages or cell types. As would be expected for the cardiac phenotype of PR interval, we found enrichment of signal in cardiac DHSs compared with DHSs from other tissue types (**Supplementary Figure 5)**. Interestingly, the second most enriched tissue DHSs were in those that regulate microvascular endothelial cells, complementing our findings (noted above) that there is enrichment in genes involved in blood vessel morphogenesis.

Using a candidate region approach in which we limited the regions examined only to those that contain cardiac DHSs (n=122,278), we identified an additional four loci associated with PR interval after Bonferroni correction for the number of SNPs tested (Table 2).

**Table 2:**
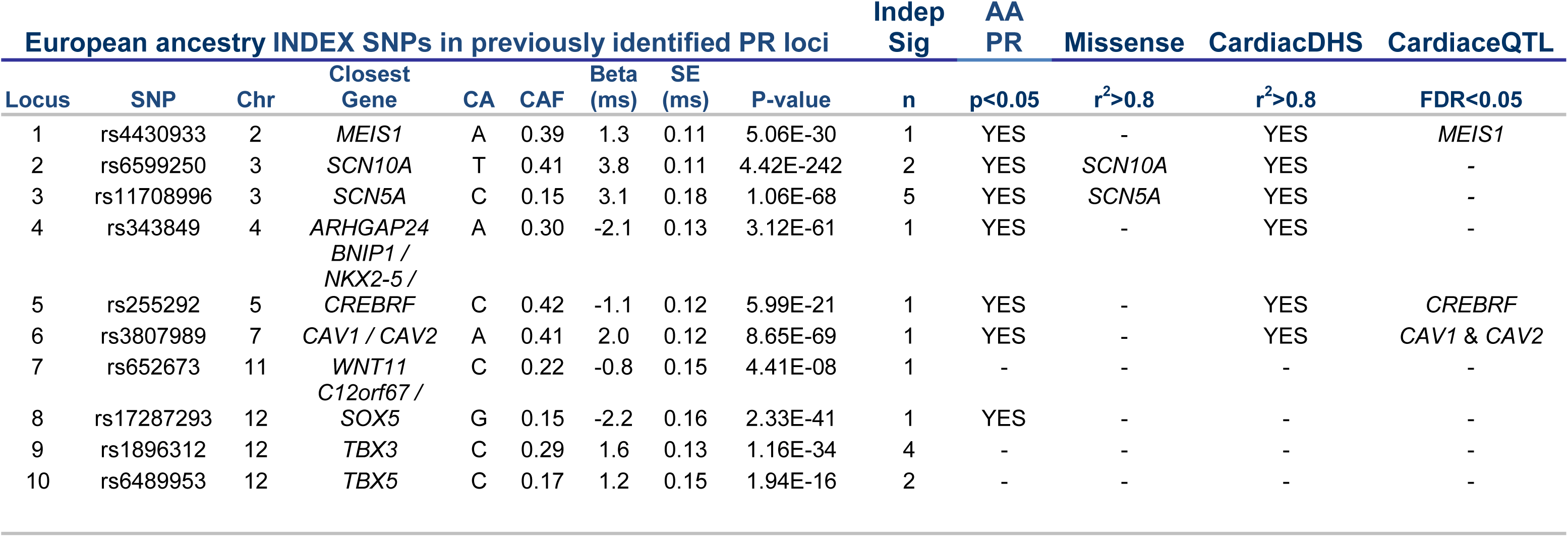
Novel loci identified by transethnic and pleiotropic meta-analyses. We combined GWAS results of Europeans and African Americans and identified an additional five loci associated with PR interval. To identify loci associated with atrioventricular conduction, we combined data on PR interval with association results of QRS duration (six novel loci), of RR interval (two novel loci), and of atrial fibrillation (three loci). Furthermore, we tested SNPs in DHSs only, adjusting the significance threshold accordingly, and found another four SNPs significantly associated with PR interval. Because some of the loci overlapped, these analyses led to 13 novel loci in total.

### Molecular Function and Biologic Processes associated with PR genes

Although extensive LD among common variants and the incompleteness of the HapMap reference panel preclude an unambiguous identification of the functional variant or the culprit gene, we used the following criteria to implicate genes in 37 of the 44 loci: (1) the gene selected was the only nearby gene (within a ±500kb window); (2) the gene selected has a missense variant in high LD (r^2^ > 0.8) with the index SNP; or (3) the index SNP was associated with cardiac transcript expression levels of the selected gene (Table 1). The set of implicated genes, detailed in **Box 1**, showed strong enrichment in genes involved in cardiac development (*P* = 1.33 × 10^−15^), specifically the cardiac chambers (*P* = 2.2 × 10^−11^) and bundle of His (*P* = 6.69 × 10^−11^) (**Supplementary Table 2**). Other notable biological processes include blood vessel morphogenesis (*P* = 7.32 × 10^−9^) and cardiac cell differentiation (*P* = 1.79 × 10^−9^). The molecular function and cellular component of the identified genes were largely enriched for transcription factors (*P* = 2.17 × 10^−6^), ion-channel related genes (*P* = 1.02 × 10^−5^), cell junction / cell signaling proteins (*P* = 4.40 × 10^−6^), and sarcomeric proteins (*P* = 4.59 × 10^−5^).

### Clinical Relevance of PR-associated Loci

To examine the clinical relevance of the identified loci, we intersected the PR genes with gene membership from multiple knowledge bases encompassing over 4,000 human diseases. The most highly over-represented conditions (*P* ≤ 1 × 10^−8^) are heart diseases including congenital abnormalities and heart failure, sick sinus syndrome and sinus arrhythmia (phenotypes related to the sinus node which houses the pacemaker cells that generate heart beats), heart block (related to cardiac conduction between atria and ventricles), and atrial fibrillation (**Supplementary Table 5**). To further explore the molecular underpinnings of these clinical relationships, we jointly analyzed the PR GWAS results with the GWAS results of heart rate^9^, QRS interval (measure of ventricular conduction)^10^, and atrial fibrillation^11^.

We examined PR SNPs for association with QRS, atrial fibrillation, and heart rate. All 61 independent SNPs from 44 loci were examined. Over half of the independent SNPs (31/61) representing 20 loci were also associated with at least one of the other electrical phenotypes (**Supplementary Table 6**, Figure 3). The cardiac sodium channel genes, *SCN5A* and *SCN10A,* clearly play a critical role in cardiac electrophysiology. PR prolonging variants in these genes are also associated with prolonged QRS duration, but with lower risks for atrial fibrillation and lower heart rate (Figure 3). The role of transcription factors in cardiac electrophysiology is equally complex. Several T-box containing transcription factors, important for cardiac conduction system formation in the developing heart, are associated with PR interval. Although *TBX3* and *TBX5* sit close together on chromosome 12, the PR prolonging allele in *TBX5* prolongs QRS and decreases AF risk while the PR prolonging allele in *TBX3* shortens QRS duration while also decreasing AF risk. The PR prolonging variant near *TBX20* prolongs QRS duration but is not associated with AF risk (Figure 3). Overall, eight of the 13 transcription factor genes associated with PR interval were also associated with at least one other atrial or atrioventricular electrical phenotype.

**Figure 3:**
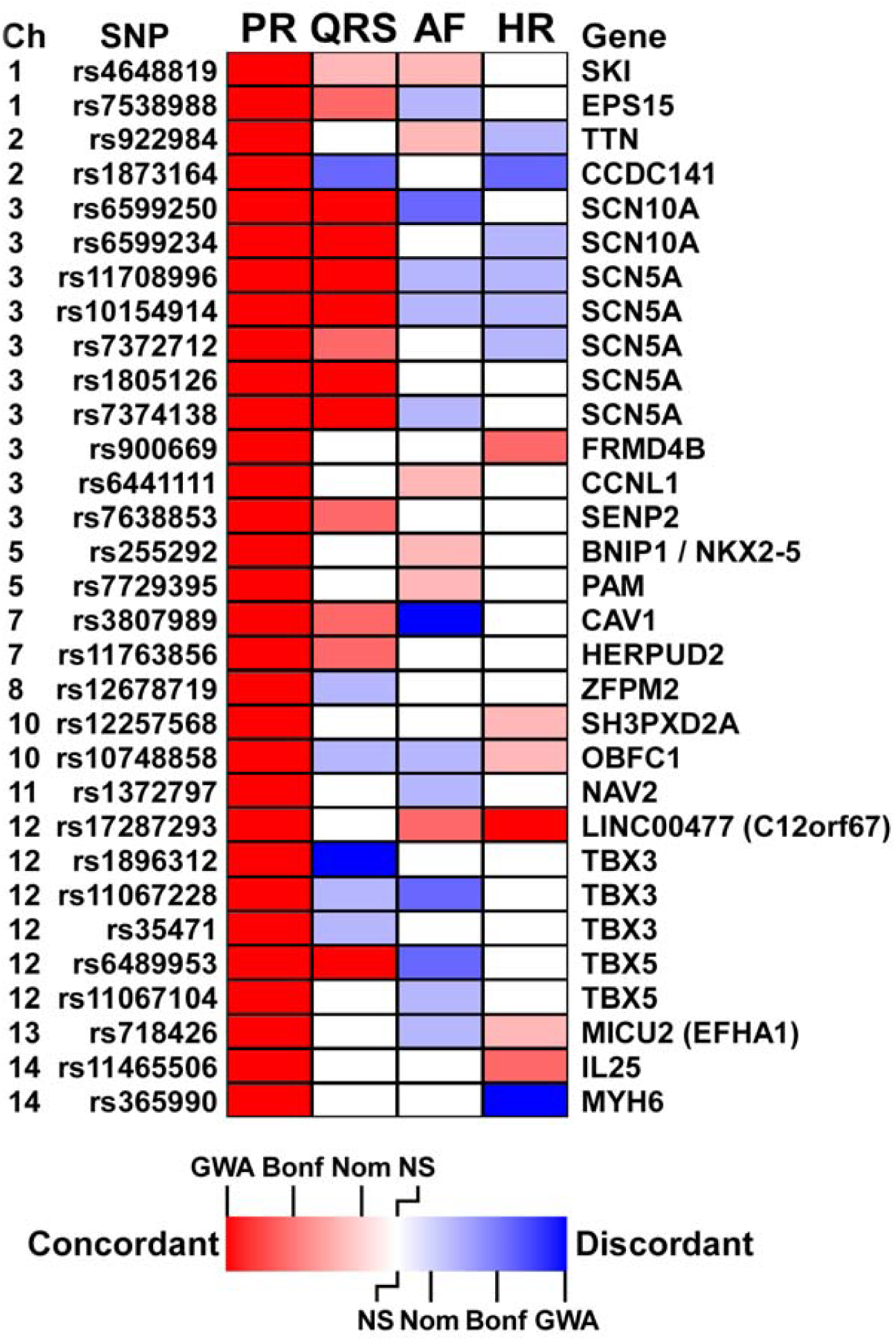
Heatmap showing overlapping loci between four traits. For each locus associated with PR interval, we tested strength of the association and direction of effect for three related traits: QRS duration, atrial fibrillation, and heart rate. While the genetic bases of these three traits show a distinct overlap with that of PR interval, we observe for each trait overlapping loci with both concordant and discordant associations, with some variants that prolong PR interval prolonging QRS duration or RR interval (concordant associations), whereas others shorten QRS duration or decrease RR interval. Similarly, some variants that prolong PR interval increase AF risk (concordant association) while others decrease AF risk (discordant).

### PR and QRS intervals

Many loci regulate both atrial / atrioventricular (PR interval) and ventricular (QRS) depolarization and conduction: twelve of our 44 PR loci were nominally associated with QRS duration (**Supplemental Table 6**) and, similarly, 17 of 22 previously identified QRS loci were at least nominally associated with PR interval (**Supplementary Table 7)**. Several intriguing findings are worth highlighting. First, while SNPs in most loci that are associated with prolonged PR are also associated with prolonged QRS, two loci have genome-wide significant discordant PR – QRS relationships, in which prolonged PR variants are associated with shorter QRS duration (*TBX3* and *SNORD56B)*; **Supplementary Table 6**, Figure 3, **Supplementary Figure 6b**. Second, although *TBX20* plays a crucial role in the development of the cardiac conduction system, the SNPs that are associated with atrial and atrioventricular conduction (PR) differ from those related to ventricular conduction (QRS) (index SNP PR rs11763856, index SNP QRS rs1419856, r^2^ = 0.001). A better understanding of the influence of these specific regions on cardiac conduction will require further investigation.

### PR interval and Atrial Fibrillation

One-third (18/61) of PR index SNPs were nominally associated with AF. Of these 18 prolonged PR SNPs, six are associated with increased AF risk, whereas 12 paradoxically lowered AF risk. This observation is consistent with the relationship between PR interval and AF described in population studies, where we showed that while both short (<120 ms) and long (>200 ms) PR interval are associated with increased AF risk, short PR interval is associated with higher risk than long PR interval.^11^ For both concordant (meaning relationships where the PR prolonging variant is associated with increased AF risk) and discordant PR – AF relationships, the larger the SNP effect size for PR interval, the larger the odds ratio for association with AF (**Supplementary Figure 6a)**. The *CAV1* index SNP associated with increased PR interval and decreased AF risk reached genome-wide significance for both phenotypes. Furthermore, of 23 previously described AF GWAS loci, 11 were at least nominally associated with PR interval.^12^ Interestingly, despite adequate power to identify modest associations, several loci, including *PITX2,* the most significant AF GWAS locus, showed no association with PR interval (**Supplementary Table 7).** Therefore, these loci may have a mode of action independent of atrial and atrioventricular depolarization or conduction.

### PR interval and Heart Rate

Ten PR loci were nominally associated with heart rate, including two sarcomeric proteins, *MYH6* and *TTN*. At the *MYH6* locus, variant rs365990 is associated prolonged PR interval and with slower heart rates, whereas an independent *MYH6* signal (<20 kb away; rs11465506) also associates with prolonged PR but is associated with markedly faster heart rates. We then examined heart rate SNPs for association with PR and found half of the heart rate SNPs were associated with PR interval, with both concordant and discordant effects. Adjusting for heart rate in the regression model did not impact the effect size or significance of the PR-genotype associations, even though resting heart rate is modestly associated with PR interval (**Supplementary Figure 7**).

### Joint phenotype meta-analyses

Finally, we performed joint phenotype analyses, with PR-heart rate, PR-QRS, and PR-atrial fibrillation as outcomes, to increase the power of finding loci involved in cardiac electrical activity. As described above, prolonged PR variants can have either a concordant or discordant association with another electrical phenotype. Therefore, we modeled the outcome for each joint analysis in two ways: with a variant having a concordant effect on PR-QRS, PR-HR, and PR-AF, and a discordant effect (**Supplementary Figures 6a-c)**. These analyses yielded 10 novel loci associated with atrial electrical activity: five related to atrial and ventricular conduction (from PR-QRS analyses); two related to atrial electrical activity and arrhythmias (from PR-AF analyses); two related to atrial depolarization and heart rate (from PR-HR analyses); and one related to both PR-QRS and PR-AF (Table 2, **Box 1, Supplementary Figure 8, Supplementary Table 8)**. Additional support for association of several of these loci were obtained by the DHS analysis, detailed above, and by trans-ethnic meta-analysis with African Americans, described below, lending further support to the validity of these associations (Table 2, **Supplementary Figure 8).**

### Trans-ethnic Analyses

Our study had less power to find associations among African Americans (n =13,415) than among European-descent individuals (n = 92,340). Nonetheless, 16 of the 44 European-identified loci nominally replicated among African Americans, suggesting that a large proportion of genetic associations with PR interval are shared between the two ethnic groups **(Supplementary Table 7)**. For European-descent GWAS PR SNPs at least nominally associated with PR among African Americans, the estimated effect was in the same direction for the two populations (**Supplementary Figure 6d)**.

Examining only the index signal may underestimate the true number of locus associations that replicate. For instance, the *TBX5* locus index SNP rs6489953 is part of a large LD block associated with PR interval among individuals of European descent. This SNP is not significantly associated with PR interval among African Americans (beta = 0.04, *P* = 0.90, **Supplementary Table 6**, Figure 2c). There is, however, a very strong SNP-PR association signal in the *TBX5* among African Americans (index SNP rs7955405, beta = 1.16, *P* = 9.2 × 10^−16^ in African Americans), Figure 2c. This SNP is in high LD with rs6489953 among European descent individuals (HapMap CEU r^2^ = 0.62), but not among populations from African descent (HapMap YRI r^2^ = 0.03). Hence, examination of only the top European descent index signal would miss the association among African Americans. Furthermore, interrogation of the *TBX5* locus among African Americans narrows the association block, allowing for fine mapping of the association signal (Figure 2c). A second noteworthy interethnic difference is that there are SNP associations among those of European descent, for instance rs1896312 in *TBX3*, where despite adequate power, no association could be established among African Americans (Figure 2c).

Our trans-ethnic GWAS meta-analysis of PR interval among 13,415 African Americans and 92,340 European-ancestry individuals identified five additional novel loci associated with atrial and atrioventricular conduction (PR interval) (Table 2, **Supplementary Figure 8**).

## Discussion

Our GWAS meta-analytic study of over 92,000 individuals of European ancestry identified 44 loci associated with cardiac atrial and atrioventricular conduction (PR interval). The implicated genes at these loci show strong enrichment for genes involved in processes related to cardiac conduction, namely, cardiovascular system development and, specifically, in development of the cardiac chamber and bundle of His. Similarly, diseases overrepresented by these genes are processes related to arrhythmias and heart block, consistent with the current knowledge of the physiology and epidemiology of cardiac atrial conduction.

Using HapMap^13^ imputation, we tested over 2.7 million SNPs, and while we did not directly test all common variants with this approach, nor did we test low-frequency variants (with minor allele frequencies below 1%), we identified many index SNPs in LD with functional variants, either through amino acid changes or involvement in gene regulation. For most newly identified loci, we are able to pinpoint a gene that may be causative, either because the index SNP (or a SNP in high LD with it) is a missense variant in the gene, or because it regulates the expression of the gene. Regulation of gene expression can be tissue specific, and our findings underscore the importance of examining eQTL data in tissue types relevant to the trait of interest.

A total of 34 novel loci were identified for PR interval in Europeans. Several have been identified previously in a related phenotype or in a different ancestral population, reassuring the validity of our results. Two loci, *EFHA1* and *LRCH1*, were previously identified for association with the PR segment.^14^ In addition, the novel locus *CAMK2D* was found to associated with P-wave duration, and *MYH6* with P-wave duration and P-wave terminal force.^15^ The *ID2* locus on chromosome 2 was found in a GWAS on PR interval in Hispanic/Latino populations.^16^ A locus that was identified in two studies in Asian populations,^17,18^ *SLC8A1*, did not reach genome-wide significance in our meta-analysis, but was suggestive with the strongest SNP being rs13026826 (beta for A-allele: 0.278, P=1.036 × 10^−6^).

Contrasting meta-analyzed association results from European descent individuals with results from a smaller sample of African Americans, we find that, with few exceptions, a large proportion of genetic associations with PR interval are shared between the two ethnic groups. We then combined data from Europeans and African Americans in a trans-ethnic meta-analysis, allowing us to find additional loci. With over 105,000 samples in total, our power to find association – even with small effect sizes – was substantial for common variants. Future studies should examine sequence or other data that provide better assessment of rare and common functional variants, as was done previously for *SCN5A*.^19^

We also combined our results on PR interval with previously published results on heart rate, QRS duration, and atrial fibrillation, and identified loci contributing to atrial arrhythmias and atrioventricular conduction. We observed significant pleiotropy of effect of these SNPs, with over half of the SNPs associated with PR interval (atrial conduction) in the study also associated ventricular conduction (QRS interval), atrial arrhythmias (atrial fibrillation), and heart rate (RR interval).

Our series of GWAS studies, including transethnic and cross-trait meta-analytic studies, has identified 57 loci, 47 of which are novel, associated with cardiac atrial and atrioventricular electrical activity among individuals of European and African ancestry. Understanding the biology of a trait in this way provides insight into related disease processes and may help identify potential approaches to drug therapy.

## References

1. Kannel, W.B. & Benjamin, E.J. Current perceptions of the epidemiology of atrial fibrillation. Cardiology clinics 27, 13–24, vii (2009).

2. Hanson, B. et al. Genetic factors in the electrocardiogram and heart rate of twins reared apart and together. The American journal of cardiology 63, 606–9 (1989).

3. Pfeufer, A. et al. Genome-wide association study of PR interval. Nature genetics 42, 153–9 (2010).

4. Holm, H. et al. Several common variants modulate heart rate, PR interval and QRS duration. Nature genetics 42, 117–22 (2010).

5. Bulik-Sullivan, B.K. et al. LD Score regression distinguishes confounding from polygenicity in genome-wide association studies. Nat Genet 47, 291–5 (2015).

6. Huang, H., Chanda, P., Alonso, A., Bader, J.S. & Arking, D.E. Gene-based tests of association. PLoS genetics 7, e1002177 (2011).

7. Kumar, P., Henikoff, S. & Ng, P.C. Predicting the effects of coding non-synonymous variants on protein function using the SIFT algorithm. Nature protocols 4, 1073–81 (2009).

8. Adzhubei, I.A. et al. A method and server for predicting damaging missense mutations. Nature methods 7, 248–9 (2010).

9. Eijgelsheim, M. et al. Genome-wide association analysis identifies multiple loci related to resting heart rate. Human molecular genetics 19, 3885–94 (2010).

10. van der Harst, P. et al. 52 Genetic Loci Influencing Myocardial Mass. J Am Coll Cardiol 68, 1435–48 (2016).

11. Ellinor, P.T. et al. Meta-analysis identifies six new susceptibility loci for atrial fibrillation. Nature genetics 44, 670–5 (2012).

12. Christophersen, I.E. et al. Large-scale analyses of common and rare variants identify 12 new loci associated with atrial fibrillation. Nat Genet 49, 946–952 (2017).

13. Frazer, K.A. et al. A second generation human haplotype map of over 3.1 million SNPs. Nature 449, 851–61 (2007).

14. Verweij, N. et al. Genetic determinants of P wave duration and PR segment. Circ Cardiovasc Genet 7, 475–81 (2014).

15. Christophersen, I.E. et al. Fifteen Genetic Loci Associated With the Electrocardiographic P Wave. Circ Cardiovasc Genet 10(2017).

16. Seyerle, A.A. et al. Genome-wide association study of PR interval in Hispanics/Latinos identifies novel locus at ID2. Heart (2017).

17. Hong, K.W. et al. Identification of three novel genetic variations associated with electrocardiographic traits (QRS duration and PR interval) in East Asians. Hum Mol Genet 23, 6659–67 (2014).

18. Sano, M. et al. Genome-wide association study of electrocardiographic parameters identifies a new association for PR interval and confirms previously reported associations. Hum Mol Genet 23, 6668–76 (2014).

19. Magnani, J.W. et al. Sequencing of SCN5A identifies rare and common variants associated with cardiac conduction: Cohorts for Heart and Aging Research in Genomic Epidemiology (CHARGE) Consortium. Circ Cardiovasc Genet 7, 365–73 (2014).

